# Understanding olfactory dysfunction in COVID-19: Expression of ACE2, TMPRSS2 and Furin in the nose and olfactory bulb in human and mice

**DOI:** 10.1101/2020.05.15.097352

**Authors:** Rumi Ueha, Kenji Kondo, Ryoji Kagoya, Shigeyuki Shichino, Satoshi Ueha, Tatsuya Yamasoba

**Author notes:** Correspondence to: Rumi Ueha, Department of Otolaryngology and Head and Neck Surgery, Faculty of Medicine, the University of Tokyo, Tokyo, Japan, 113-8655.

## Abstract

**Background:** Anosmia is a frequent symptom in coronavirus disease 2019 (COVID-19) patients that generally resolves within weeks. In contrast, the anosmia caused by other upper respiratory infections affects a small proportion of patients and may take months to resolve or never resolve. The mechanisms behind COVID-19-induced olfactory dysfunction remain unknown. Here, we address the unique pathophysiology of COVID-19-associated olfactory dysfunction.

**Methods:** The expression of ACE2 (virus binding receptor) and TMPRSS2 and Furin (host cell proteases facilitating virus entry) was examined in the nasal mucosa, composed of respiratory mucosa (RM), olfactory mucosa (OM), and olfactory bulb (OB) of mouse and human tissues using immunohistochemistry and gene analyses.

**Results:** Co-expression of ACE2, TMPRSS2, and Furin was observed in the RM and in the OM, especially in the supporting cells of the olfactory epithelium and the Bowman’s glands. Notably, the olfactory receptor neurons (ORNs) in the OM were positive for ACE2 but almost negative for TMPRSS2 and Furin. Cells in the OB expressed ACE2 strongly and Furin weakly, and did not express TMPRSS2. All three gene expressions were confirmed in the nasal mucosa and OB.

**Conclusions:** ACE2 was widely expressed in all tissues, whereas TMPRSS2 and Furin were expressed only in certain types of cells and were absent in the ORNs. These findings, together with clinical reports, suggest that COVID-19-related anosmia occurs mainly through sensorineural and central dysfunction and, to some extent, conductive olfactory dysfunction. The expression of ACE2, but not TMPRSS2 or Furin, in ORNs may explain the early recovery from anosmia.

## Introduction

The recent international spread of the coronavirus disease 2019 (COVID-19) caused by severe acute respiratory syndrome coronavirus 2 (SARS-CoV-2) poses a serious health emergency. COVID-19 usually begins with simple respiratory symptoms such as fever, shore throat, and cough for 2-3 days (*1, 2*). Notably, chemosensitive disorders, such as loss or decline of smell (anosmia or hyposmia), and loss of taste (ageusia or dysgeusia), have been repeatedly reported as unique clinical features of COVID-19 (*3, 4*), and are now considered typical symptoms of the early stages of SARS-CoV-2 infection (*4, 5*). A recent published meta-analysis demonstrated a 52.73% prevalence of olfactory dysfunction among 1,627 COVID-19 patients (*6*). Specifically, a high rate of anosmia (complete loss of smell) is well documented (*3, 5, 7, 8*). In contrast, only a small proportion (up to 20%) of patients with upper respiratory infection (URI) exhibits olfactory dysfunction (*9*). The prognosis of olfactory dysfunction due to URI is generally poor, with the majority of patients showing no or slight recovery within a few months (*10*), whereas olfactory dysfunction in COVID-19 patients resolves relatively rapidly, with a reported duration of 1 to 4 weeks (*3, 8, 11-13*). In a cohort study of COVID-19 patients with olfactory dysfunction, loss of smell was reported to be the first symptom in 27% of patients (*12*). Thus, the clinical features of COVID-19-associated olfactory dysfunction are significantly different from those found in patients with common URIs, which are caused by viruses such as rhinovirus, picornavirus, and parainfluenza virus (*9, 14-16*).

SARS-CoV-2 host cell entry depends on several factors: the binding of viral spike proteins to the cellular receptor angiotensin-converting enzyme 2 (ACE2) (*17-19*), spike protein cleavage by the host cell enzyme Furin (*19-21*), and spike protein priming by host cell proteases such as transmembrane protease serine 2 (TMPRSS2) (*19, 22*). Thus, the high expression of ACE2, TMPRSS2, and Furin is thought to enhance SARS-CoV-2 entry as well as clinical symptoms.

Based on its histological components and functions, the nasal mucosa is divided into the respiratory mucosa (RM) and the olfactory mucosa (OM). The RM consists of various types of epithelial cells, including ciliated columnar and goblet cells. The OM serves olfaction and consists of the olfactory epithelium (OE) and subepithelial tissues (*23*). The degree of olfaction is closely related to the number of mature olfactory receptor neurons (ORNs) in the OE. The olfactory system consists of peripheral compartments such as the OM, and central structures such as the olfactory bulb (OB) and the piriform/entorhinal cortex (*24*).

To date, the expression of *ACE2* (*25, 26*) and *TMPRSS2* (*26*), but not *Furin*, has been reported in the nasal epithelium. However, the histological evaluation of their expression has been limited; only one study reported the expression of ACE2 and TMPRSS2 proteins in the nasal mucosa and respiratory sinus, although it did not demonstrate immunostaining images of the nasal mucosa (*27*). In the present study, we sought to elucidate the mechanisms underlying olfactory dysfunction and the pathogenesis of high viral load in the upper airways of COVID-19 patients. Toward that aim, we investigated the expression of ACE2, TMPRSS2, and Furin in the RM, OM, and OB of human and mouse tissues.

## Methods

### Experimental samples

Animal tissue samples were all obtained from the mice examined in the previous published studies (*23, 28*), because purchasing new animals had been prohibited in our facility due to the epidemic spread. The samples from 6 eight-week-old male C57BL/6 mice (*23*) and an eight-week-old-male Sprague Dawley rat (*28*) were used, and the following paraffin-embedded tissues were collected; the RM area and OM area of the nose, the OB area, and the kidney and prostate for positive controls of immunostaining (Figure 1A). Human tissues were obtained from patients undergoing surgery for the treatment of chronic sinusitis or olfactory neuroblastoma. These included the OM (n = 3), the middle turbinate (n = 5), and inferior turbinate (n = 6); the latter two were used for the evaluation of the RM. Routine morphology was evaluated in haematoxylin and eosin-stained sections by a qualified pathologist and otolaryngologists. Tissue evaluation was performed only in the parts characterized as non-diseased. All experiments were conducted in accordance with institutional guidelines and with the approval of the Animal Care and Use Committee of the University of Tokyo (No. P14-051, P15-115) and of the Research Ethics Committee of the Graduate School of Medicine and Faculty of Medicine, the University of Tokyo, Japan (12009, 2019073NI). Since archived specimens were used, written informed consent was waived.

**Figure 1.**
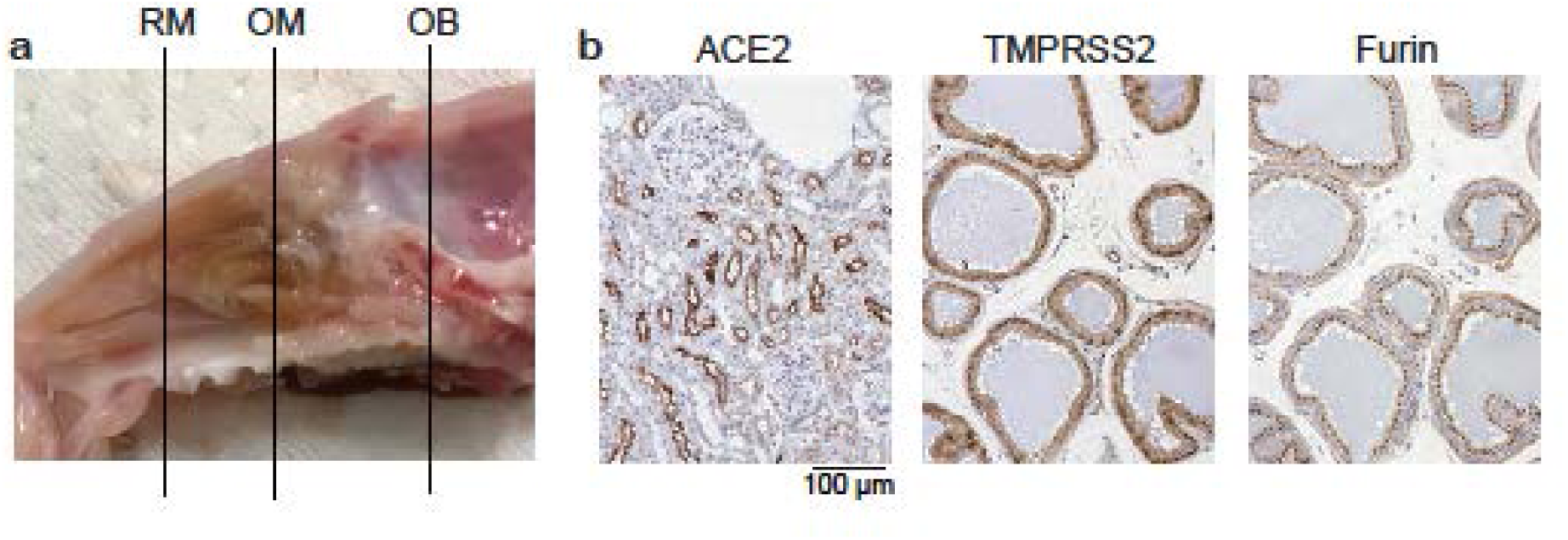
Nose sagittal section and representative images of ACE2 staining in mouse kidney, and TMPRSS2 and Furin staining in rat prostate. **a**: Sagittal section of the rodent nose. The bars indicate the respiratory mucosal section (RM), olfactory mucosal section (OM), and the olfactory bulb section (OB). **b**: Positive staining for ACE2 in the kidney, and TMPRSS2 and Furin in the prostate are shown. ACE2 positive cells are observed in the brush boarder and the cytoplasm and nucleus of tubular cells. TMPRSS2 positive cells are observed in the cytoplasm and nucleus of acinar cells. Furin positive cells are observed in the supranuclear cytoplasm (mainly Golgi apparatus) of acinar cells in the prostate glands.

### Histological analyses

To detect the expressions of ACE2 and TMPRSS2 in the RM, OM, and OB, histological analyses were performed by immunostaining. Four-µm-thick serial paraffin sections were deparaffinized in xylene and dehydrated in ethanol before immunostaining. Prior to immunostaining, deparaffinized sections were treated with 3% hydrogen peroxide to block endogenous peroxidase activity and were incubated in Blocking One (Nacalai Tesque, Kyoto, Japan) to block non-specific immunoglobulin binding. After antigen activation, primary antibodies against ACE2 (1:300 dilution; rabbit monoclonal, Abcam, ab108252; Cambridge, UK), TMPRSS2 (1:1000 dilution; rabbit monoclonal, Abcam, ab92323; Cambridge, UK), Furin (1:100 dilution; rabbit monoclonal, Abcam, ab183495; Cambridge, UK), and PGP9.5 for a neuronal marker (1:500 dilution; guinea pig polyclonal, Abcam, ab10410; Cambridge, UK) were detected with peroxidase conjugated appropriate secondary antibodies and a diaminobenzidine substrate. The mouse kidney and rat prostate were stained for positive controls for ACE2 and for TMPRSS2 and Furin, respectively (Figure 1B). All samples were stained under the same condition and protocol as the positive control staining. Images of all sections were captured using a digital microscope camera (Keyence BZ-X700) with 4×, 10x, 20×, and 40x objective lenses.

### Gene expression analyses

Our previous DNA microarray data from the nasal mucosa and OB (NCBI Gene Expression Omnibus database under the series number GSE 103191, 150694) was used to examine the expressions of *ACE2, TMPRSS2*, and *Furin*. The expression levels of each gene were normalized against the expression level of *Rps3* (encoding ribosomal protein S3) in each sample.

## Results

The immunohistological data is summarized in Table 1. ACE2, TMPRSS2, and Furin were present in human and mouse nasal mucosa and in mouse OB, though the expression pattern of ACE2, TMPRSS2, and Furin varied among tissues. Remarkably, co-expression of ACE2, TMPRSS, and Furin was detected in the supporting cells and Bowman’s glands of the OM, and diffusely in the RM, but not in the ORNs of the OM or in the OB.

**Table 1.** Protein expression levels of ACE2, TMPRSS2, and Furin in the respiratory mucosa, olfactory mucosa, and olfactory bulb. Both the respiratory mucosa (RM) and olfactory mucosa (OM) can be further subdivided into the epithelium and the lamina propia (LP). In the respiratory epithelium (RE) several cells such as ciliated columnar and globet cells can be found, while the subepithelial glands are found in the LP. The OM, which serves olfaction, consists of an olfactory epithelium (OE), which is composed of olfactory receptor neurons (ORNs) and supporting and basal cells, and the LP, which contains Bowman’s glands and olfactory nerve bundles. Importantly, the degree of olfaction is closely related to the number of mature olfactory receptor neurons (ORNs) in the OE. Moreover, the olfactory system consists of peripheral compartments such as the OM, and central structures such as the olfactory bulb (OB) and the piriform/entorhinal cortex. The olfactory bulb consists of three layers; the glomerular, mitral cell, and granule cell layers. +: mild expression, ++: moderate to strong expression, -: negative.

|  |  |  | ACE2 | TMPRSS2 | Furin |
| --- | --- | --- | --- | --- | --- |
| RM | RE | Nucleus | ++ | + | - |
|  |  | Cytoplasm | + | + | + |
|  | LP | Subepithelial Gland | + | ++ | ++ |
| OM | OE | Supporting cell | ++ | + | + |
|  |  | Olfactory receptor neuron | ++ | - | - |
|  |  | Basal cell | ++ | ± | - |
|  |  | Cytoplasm | - | + | + |
|  | LP | Bowman's Gland | + | ++ | ++ |
|  |  | Olfactory nerve bundle | + | + | - |
| OB | Glomerular layer |  | ++ | - | + |
|  | Mitral cell layer |  | ++ | - | + |
|  | Granule cell layer |  | ++ | - | - |

In mouse RM, ACE2, TMPRSS2, and Furin were all strongly expressed in the cytoplasm of respiratory epithelial cells and in the subepithelial glands (Fig. 1a, b). ACE2 and TMPRSS2 were highly co-expressed in the RE. The villous brush border of the respiratory columnar epithelium was strongly positive for TMPRSS2 expression. In addition, moderate cytoplasmic staining for TMPRSS2 and Furin was observed in the subepithelial tissue (Fig. 2a, b). In OM, only supporting cells and Bowman’s glands expressed ACE2, TMPRSS2, and Furin. The olfactory nerve bundles were moderately positive for ACE2 and TMPRSS2. Notably, all cells in the OE, including supporting cells, ORNs, and basal cells, were positive for ACE2, while the ORNs were negative for TMPRSS2 and Furin (Fig. 2c, d). In the OB, there were no cells expressing ACE2, TMPRSS2, and Furin simultaneously (Fig. 3a-c). ACE2 positive cells were detected in the glomerular layer, mitral cell layer, and granule cell layer, but those cells were negative for TMPRSS2. On the other hand, some cells in the glomerular layer and mitral cells were positive for Furin. TMPRSS2 was strongly expressed in cells of the OB core (Fig. 3b).

**Figure 2:**
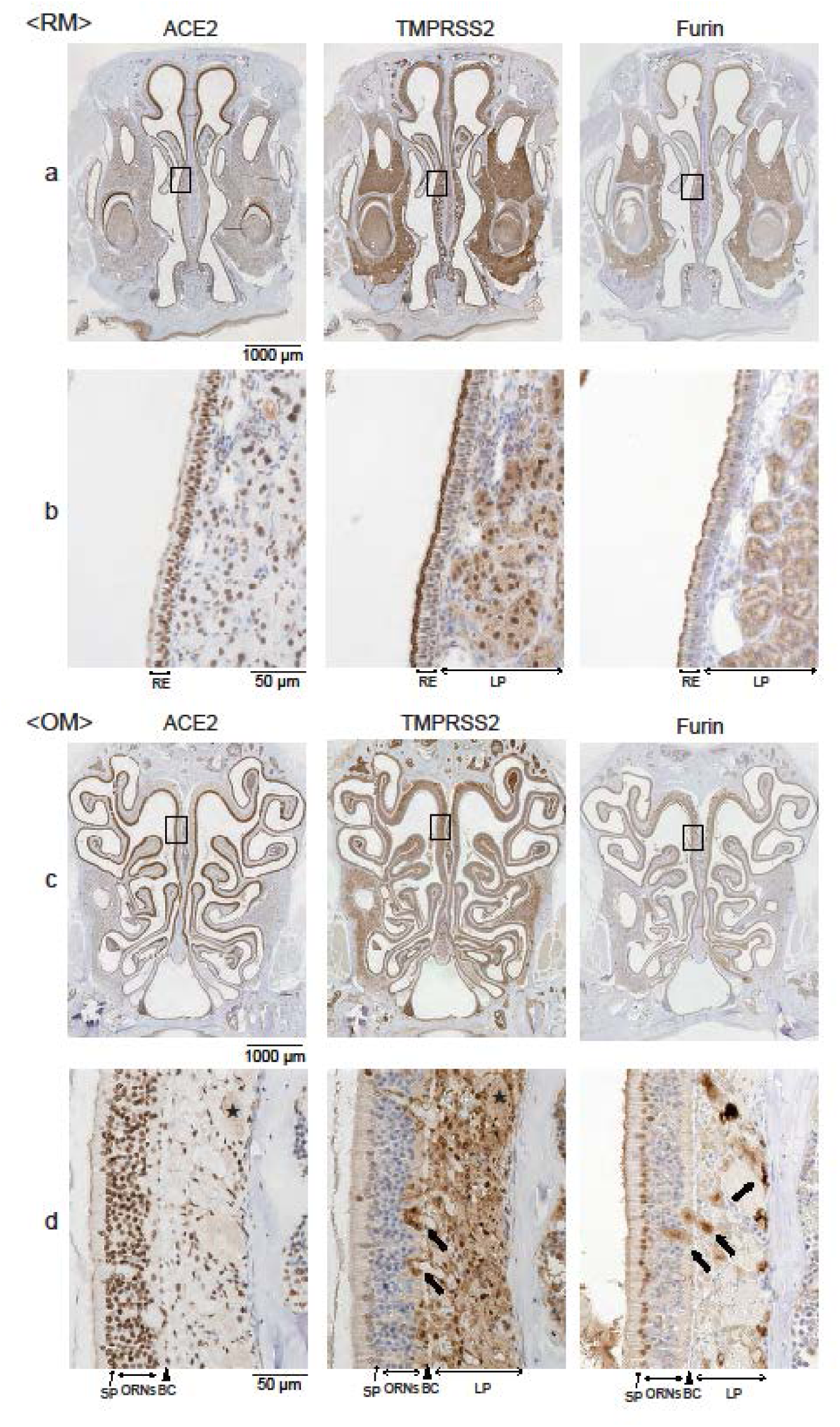
Representative images of mouse respiratory and olfactory mucosa stained with antibodies against ACE2, TNPRSS2, and Furin. **a** and **b**: The boxes in (a) indicate the parts of the respiratory mucosa (RM) that are shown at higher magnification in (b) (a, 40x magnification; b, 400x magnification). In the RM area, the respiratory epithelium (RE) was stained with ACE2 antibody. On the other hand, TMPRSS2 and Furin were strongly stained in the cilia of the ciliated columnar RE and moderately stained in the supranuclear cytoplasm of RE cells and subepithelial glands. **c** and **d**: In the olfactory mucosal (OM) area, supporting cells (SPs, arrows), olfactory receptor neurons (ORNs), and basal cells (BCs, arrow heads) were strongly stained with ACE2 antibody. TMPRSS2 expression was negative in ORNs, moderately positive in SPs, and strongly positive in Bowman’s glands (thick black arrows) and the lamina propria. The olfactory nerve bundles showed weak co-expression of ACE2 and TMPRSS2 (star marks). Furin was also expressed in SPs and Bowman’s glands (thick black arrows) The boxes in (c) indicate the parts of the OM that are shown at higher magnification in (d) (c, 40x magnification; d, 400x magnification).

**Figure 3:**
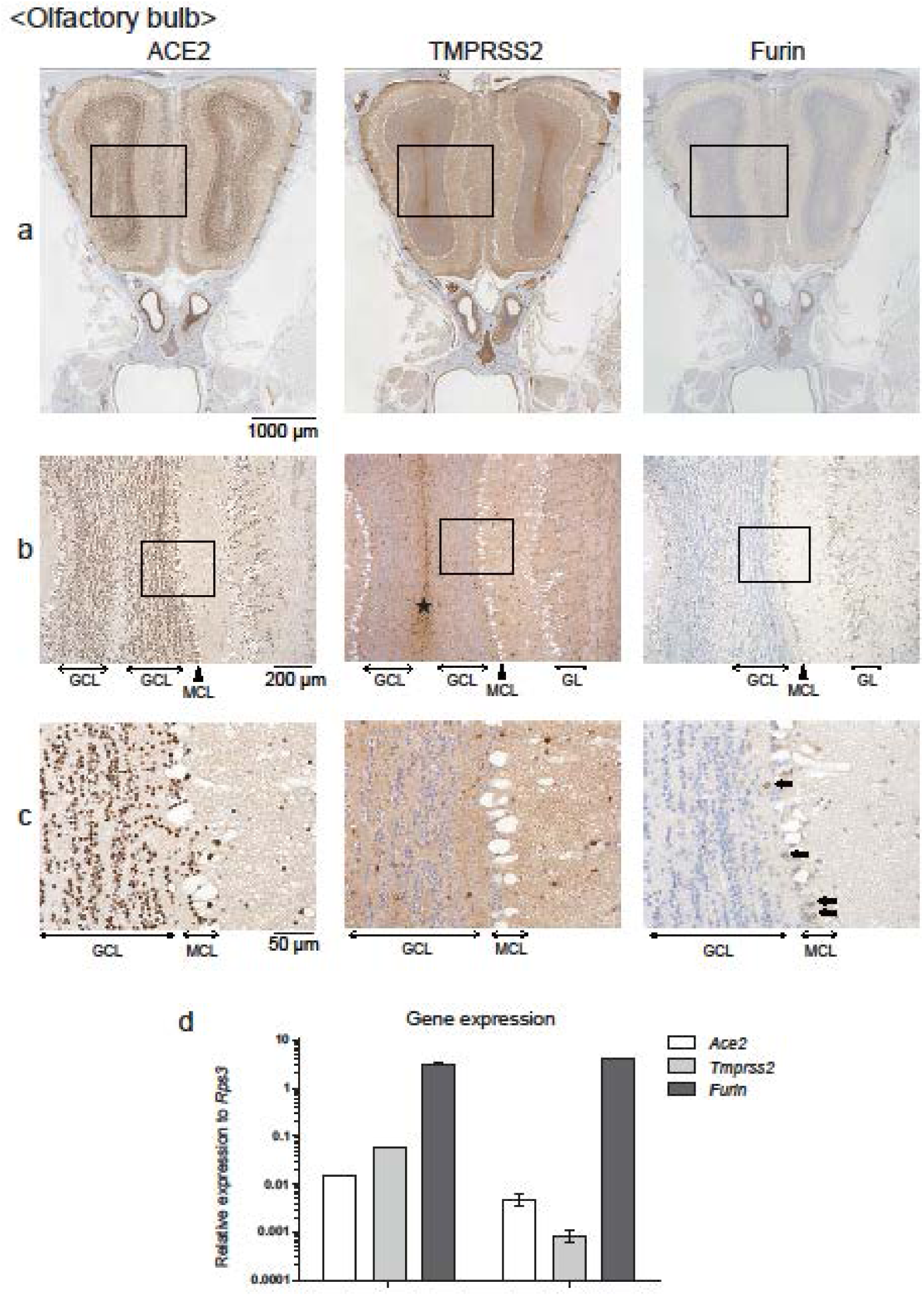
Representative images of mouse olfactory bulb stained with antibodies against ACE2, TNPRSS2, and Furin; and gene expression levels in the nasal mucosa and olfactory bulb. **a, b, c**: The olfactory bulb (OB) area. The boxes in (a) and (b) are shown at higher magnification in (b) and (c), respectively (a, 40x magnification; b, 100x magnification; c, 400x magnification). The OB parenchyma was weakly positive for ACE2, strongly positive for TMPRSS2, and negative for Furin. While ACE2 positive cells could be recognized in the glomerular layer (GL), mitral cell layer (MCL, arrow heads), and granule cell layer (GCL), those cells were negative for TMPRSS2. Some mitral cells were positive for Furin (thick black arrows). TMPRSS2 was strongly expressed in the cells of the OB core (star marks). **d**: Gene expression levels of *Ace2, Tmprss2*, and *Furin* in the nasal mucosa and olfactory bulb are shown relative to the expression of the endogenous control gene *Rps3* (encoding ribosomal protein S3). The graph is shown in log_10_ scale.

To reinforce the above histological results, we investigated *Ace2, Tmprss2*, and *Furin* gene expression in mouse nasal mucosa and OB. Using the database from the previous study(*24*), the expression of the three genes was confirmed in the nasal mucosa and OB (Fig. 3d).

In human nasal mucosa, the PGP9.5 antibody clearly visualized the OE containing olfactory neurons. ACE2 was localized in PGP9.5 positive ORNs. In addition, while Furin was not present in the OE, TMPRSS2 was weakly expressed in the apical layer of the OE. (Fig. 4a, b). In the RM, ACE2, TMPRSS2, and Furin were widely co-expressed in the epithelium (Fig. 4c, d). These findings were basically identical to those found in mouse tissues.

**Figure 4:**
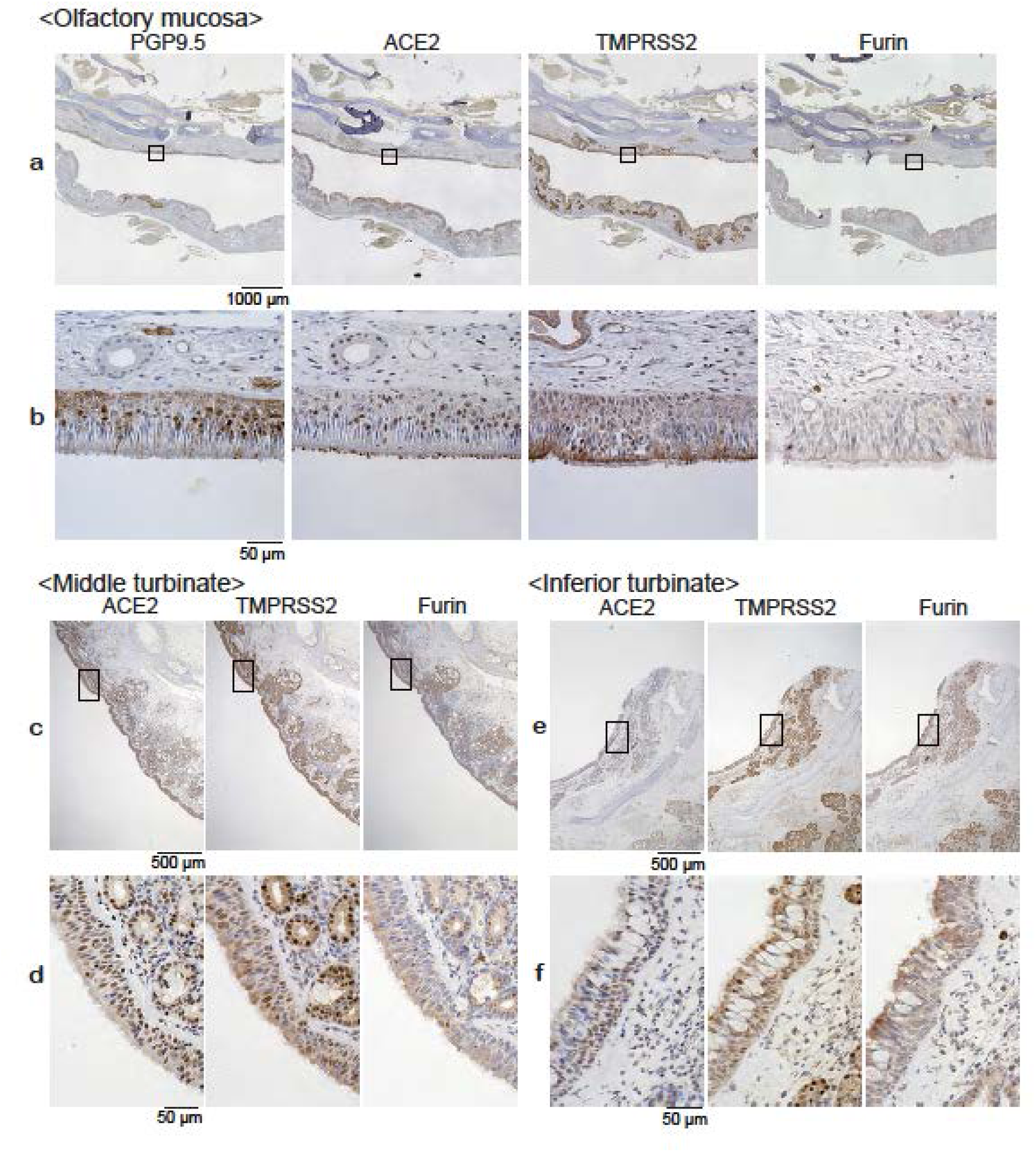
Representative images of human tissues stained with antibodies against ACE2, TMPRSS2, and Furin. **a** and **b**: In the olfactory mucosal area, the area of the olfactory epithelium was recognized using PGP9.5 staining. PGP9.5^+^ ORNs mostly expressed ACE2, but not TMPRSS2 or Furin. There was high expression of TMPRSS2 in the cytoplasm of the subepithelial glands. Furin was rarely detected in human olfactory epithelium, but a weak expression was noted in the subepithelial glands. The boxes in (a) are shown at higher magnification in (b) (a, 40x magnification; b, 400x magnification). **c** - **f**: Representative images of the middle turbinate and inferior turbinate. The boxes in (c) and (e) indicate the parts of the respiratory mucosa that are shown at higher magnification in (d) and (f) (c, e, 40x magnification; d, f, 400x magnification). ACE2, TMPRSS2 and Furin protein expression was detected in the respiratory epithelium and subepithelial glands. Specifically, TMPRSS2 expression was well detected.

## Discussion

COVID-19 causes numerous clinical symptoms, including a deteriorated sense of taste and smell, respiratory, and digestive disorders (*3, 4, 29*). Anosmia, the loss of smell, is a unique clinical feature observed in the early stages of COVID-19. Unlike the anosmia caused in other URIs, the COVID-19 anosmia occurs in most patients and resolves rather quickly. SARS-CoV-2 cell entry is dependent on the expression of the host cell proteins ACE2, TMPRSS2, and Furin.

Since the expression of these proteins in the nasal mucosa remains rather elusive, we investigated the expression of ACE2, TMPRSS2, and Furin in the RM, OM, and OB of human and mouse tissues to elucidate the mechanisms underlying olfactory dysfunction and high viral load in the upper airways of COVID-19 patients.

The results of the present study explain why olfaction is frequently impaired in COVID-19 patients. We observed the immunolocalization of ACE2, TMPRSS2, and Furin in the nasal tissue and the OB, which are considered to play a pivotal role in the manifestation of the olfactory dysfunction induced by SARS-CoV-2 infection. Histologically, ACE2, TMPRSS, and Furin were co-expressed in the supporting cells and Bowman’s glands of the OM as well as in the RM, but not in the ORNs of the OM or the OB. *Ace2, Tmprss2*, and *Furin* gene expression was confirmed in the nasal mucosa and the OB, supporting the immunohistochemical findings.

Olfactory dysfunction is defined into three types according to the anatomical location; conductive, sensorineural, and central (*30*). Conductive dysfunction results from the blockage of odorant airflow to the OE. Sensorineural dysfunction is caused by damage of the ORNs and the olfactory nerve, impaired olfactory adaptation, and/or odorant transport. The supporting cells and the Bowman’s glands are involved in olfactory adaptation and the neurotrophic and physical support of the OE (*31-33*). Thus, if the function of the supporting cells and the Bowman’s glands is deteriorated, odor adaptation is impaired, and subsequently, sensorineural dysfunction occurs. Central dysfunction occurs with the damage of olfactory processing pathways in the central nervous system (*30*). Based on the present histological results, we suggest that SARS-CoV-2 mainly induces sensorineural olfactory dysfunction without olfactory neuronal damage.

In the olfactory mucosa, co-expression of ACE2, TMPRSS2, and Furin in the supporting cells and the Bowman’s glands suggests that COVID-19 may induce the deterioration of mucus production and OE support, resulting in impaired odor adaptation and transduction. Moreover, co-expression of ACE2 and TMPRSS2 in the olfactory nerve bundle implies that odor transduction may be impaired through neuronal dysfunction. It is unlikely that SARS-CoV-2 directly damages the ORNs in the OM because the ORNs expressed ACE2, but not TMPRSS2 or Furin. Furthermore, the severe damage of ORNs would not explain the early recovery of olfaction observed in COVID-19 patients with anosmia, since the turnover rate of ORNs is approximately 30 days (*34*). The co-expression of ACE2 and Furin in the mitral cells of the OB, which have large cell bodies and secondary dendrites, suggests that central olfactory dysfunction may occur as a result of synaptic inhibition from the ORNs to the olfactory bulb. Although most COVID-19 patients recover from olfactory dysfunction, some patients have not recovered their olfaction after several months (*8*). Those COVID-19 patients with prolonged olfactory dysfunction may have suffered from continuing sensorineural and central olfactory dysfunction.

Considering the high expression of ACE2, TMPRSS2, and Furin in the RE and the subepithelial glands, SARS-CoV-2 possibly induces conductive olfactory dysfunction through hypersecretion and goblet cell hyperplasia (*14, 35*). In fact, a significant number of COVID-19 patients suffer from nasal obstruction and rhinorrhea(*3*), but the olfactory symptoms due to conductive olfactory dysfunction may fluctuate and mostly resolve completely. In addition, considering that a certain proportion of COVID-19 patients with hyposmia and anosmia did not exhibit nasal obstruction or rhinitis symptoms(*3, 36*), the contribution of conductive olfactory loss may be limited. However, some COVID-19 patients with prolonged olfactory dysfunction possibly suffer from continuing sensorineural and central olfactory dysfunction. Future studies are needed for clinical evaluation with long-term-follow-up. As for limitations of the present study, the evaluation was performed on mice and some patients with olfactory neuroblastoma which may not correctly reflect the backgrounds of SARS-CoV-2 infection. Future clinical case studies and autopsy studies might strengthen this study.

In conclusion, the present study demonstrates high expression of ACE2, TMPRSS2, and Furin in the nasal mucosa including the SPs and Bowman’s glands, and in OB. The expression pattern suggests that COVID-19 associated olfactory dysfunction is mainly sensorineural and central, and to some extent, conductive. The expression of ACE2, but not TMPRSS2 or Furin, in ORNs may explain the early recovery from anosmia.

## Author contributions

R.U. developed the concept, designed and performed the experiments, and wrote the draft of the manuscript. R.K. and S.S. designed and performed the experiments. S.U. performed the experiments and wrote partially the draft of the manuscript. K.K, and T.Y. developed the concept, designed the experiments and reviewed the manuscript. All authors contributed to interpretation of the data and writing of the manuscript.

## Data and materials availability

The datasets generated for this study are available on request to the corresponding author.

## References

1. C. Huang et al., Clinical features of patients infected with 2019 novel coronavirus in Wuhan, China. Lancet 395, 497 (Feb 15, 2020).

2. X. Huang, F. Wei, L. Hu, L. Wen, K. Chen, Epidemiology and Clinical Characteristics of COVID-19. Archives of Iranian medicine 23, 268 (Apr 1, 2020).

3. J. R. Lechien et al., Olfactory and gustatory dysfunctions as a clinical presentation of mild-to-moderate forms of the coronavirus disease (COVID-19): a multicenter European study. European archives of oto-rhino-laryngology: official journal of the European Federation of Oto-Rhino-Laryngological Societies, (Apr 6, 2020).

4. L. A. Vaira et al., Objective evaluation of anosmia and ageusia in COVID-19 patients: Single-center experience on 72 cases. Head & neck, (Apr 27, 2020).

5. J. Krajewska, W. Krajewski, K. Zub, T. Zatonski, COVID-19 in otolaryngologist practice: a review of current knowledge. European archives of oto-rhino-laryngology: official journal of the European Federation of Oto-Rhino-Laryngological Societies, (Apr 18, 2020).

6. J. Y. Tong, A. Wong, D. Zhu, J. H. Fastenberg, T. Tham, The Prevalence of Olfactory and Gustatory Dysfunction in COVID-19 Patients: A Systematic Review and Meta-analysis. Otolaryngology--head and neck surgery: official journal of American Academy of Otolaryngology-Head and Neck Surgery, 194599820926473 (May 5, 2020).

7. B. Russell et al., Anosmia and ageusia are emerging as symptoms in patients with COVID-19: What does the current evidence say? Ecancermedicalscience 14, ed98 (2020).

8. C. Hopkins, P. Surda, E. Whitehead, B. N. Kumar, Early recovery following new onset anosmia during the COVID-19 pandemic - an observational cohort study. Journal of otolaryngology - head & neck surgery = Le Journal d’oto-rhino-laryngologie et de chirurgie cervico-faciale 49, 26 (May 4, 2020).

9. A. M. Seiden, Postviral olfactory loss. Otolaryngologic clinics of North America 37, 1159 (Dec, 2004).

10. M. Yamagishi, M. Fujiwara, H. Nakamura, Olfactory mucosal findings and clinical course in patients with olfactory disorders following upper respiratory viral infection. Rhinology 32, 113 (Sep, 1994).

11. C. H. Yan, F. Faraji, D. P. Prajapati, C. E. Boone, A. S. DeConde, Association of chemosensory dysfunction and Covid-19 in patients presenting with influenza-like symptoms. International forum of allergy & rhinology, (Apr 12, 2020).

12. R. Kaye, C. W. D. Chang, K. Kazahaya, J. Brereton, J. C. Denneny, 3rd, COVID-19 Anosmia Reporting Tool: Initial Findings. Otolaryngology--head and neck surgery: official journal of American Academy of Otolaryngology-Head and Neck Surgery, 194599820922992 (Apr 28, 2020).

13. A. R. Sedaghat, I. Gengler, M. M. Speth, Olfactory Dysfunction: A Highly Prevalent Symptom of COVID-19 With Public Health Significance. Otolaryngology--head and neck surgery: official journal of American Academy of Otolaryngology-Head and Neck Surgery, 194599820926464 (May 5, 2020).

14. M. Suzuki et al., Identification of viruses in patients with postviral olfactory dysfunction. The Laryngoscope 117, 272 (Feb, 2007).

15. D. A. Deems et al., Smell and taste disorders, a study of 750 patients from the University of Pennsylvania Smell and Taste Center. Archives of otolaryngology--head & neck surgery 117, 519 (May, 1991).

16. W. S. Cain, J. F. Gent, R. B. Goodspeed, G. Leonard, Evaluation of olfactory dysfunction in the Connecticut Chemosensory Clinical Research Center. The Laryngoscope 98, 83 (Jan, 1988).

17. W. Li et al., Angiotensin-converting enzyme 2 is a functional receptor for the SARS coronavirus. Nature 426, 450 (Nov 27, 2003).

18. F. Wu et al., A new coronavirus associated with human respiratory disease in China. Nature 579, 265 (Mar, 2020).

19. S. Lukassen et al., SARS-CoV-2 receptor ACE2 and TMPRSS2 are primarily expressed in bronchial transient secretory cells. The EMBO journal, e105114 (Apr 4, 2020).

20. A. C. Walls et al., Structure, Function, and Antigenicity of the SARS-CoV-2 Spike Glycoprotein. Cell 181, 281 (Apr 16, 2020).

21. B. Coutard et al., The spike glycoprotein of the new coronavirus 2019-nCoV contains a furin-like cleavage site absent in CoV of the same clade. Antiviral research 176, 104742 (Apr, 2020).

22. S. Matsuyama et al., Enhanced isolation of SARS-CoV-2 by TMPRSS2-expressing cells. Proceedings of the National Academy of Sciences of the United States of America 117, 7001 (Mar 31, 2020).

23. R. Ueha, S. Ueha, K. Kondo, H. Nishijima, T. Yamasoba, Effects of Cigarette Smoke on the Nasal Respiratory and Olfactory Mucosa in Allergic Rhinitis Mice. Frontiers in neuroscience 14, 126 (2020).

24. R. Ueha et al., Reduction of Proliferating Olfactory Cells and Low Expression of Extracellular Matrix Genes Are Hallmarks of the Aged Olfactory Mucosa. Frontiers in aging neuroscience 10, 86 (2018).

25. X. Zou et al., Single-cell RNA-seq data analysis on the receptor ACE2 expression reveals the potential risk of different human organs vulnerable to 2019-nCoV infection. Frontiers of medicine, (Mar 12, 2020).

26. W. Sungnak et al., SARS-CoV-2 entry factors are highly expressed in nasal epithelial cells together with innate immune genes. Nature medicine, (Apr 23, 2020).

27. S. Bertram et al., Influenza and SARS-coronavirus activating proteases TMPRSS2 and HAT are expressed at multiple sites in human respiratory and gastrointestinal tracts. PloS one 7, e35876 (2012).

28. R. Ueha et al., Laryngeal mucus hypersecretion is exacerbated after smoking cessation and ameliorated by glucocorticoid administration. Toxicology letters 265, 140 (Jan 4, 2017).

29. L. Pan et al., Clinical Characteristics of COVID-19 Patients With Digestive Symptoms in Hubei, China: A Descriptive, Cross-Sectional, Multicenter Study. The American journal of gastroenterology 115, 766 (May, 2020).

30. T. Hummel et al., Position paper on olfactory dysfunction. Rhinology 56, 1 (Jan 31, 2016).

31. M. Okano, S. F. Takagi, Secretion and electrogenesis of the supporting cell in the olfactory epithelium. The Journal of physiology 242, 353 (Oct, 1974).

32. B. Branigan, P. Tadi, in StatPearls. (Treasure Island (FL), 2020).

33. C. Acevedo, K. Blanchard, J. Bacigalupo, C. Vergara, Possible ATP trafficking by ATP-shuttles in the olfactory cilia and glucose transfer across the olfactory mucosa. FEBS letters 593, 601 (Mar, 2019).

34. D. G. Moulton, Dynamics of cell populations in the olfactory epithelium. Annals of the New York Academy of Sciences 237, 52 (Sep 27, 1974).

35. T. Hummel, Perspectives in Olfactory Loss Following Viral Infections of the Upper Respiratory Tract. Archives of otolaryngology--head & neck surgery 126, 802 (Jun, 2000).

36. L. A. Vaira, G. Salzano, G. Deiana, G. De Riu, Anosmia and Ageusia: Common Findings in COVID-19 Patients. The Laryngoscope, (Apr 1, 2020).

